# SEMA: Antigen B-cell conformational epitope prediction using deep transfer learning

**DOI:** 10.1101/2022.06.20.496780

**Authors:** Tatiana I. Shashkova, Dmitriy Umerenkov, Mikhail Salnikov, Pavel V. Strashnov, Alina V. Konstantinova, Ivan Lebed, Dmitrii N. Shcherbinin, Marina N. Asatryan, Olga L. Kardymon, Nikita V. Ivanisenko

**Affiliations:** AIRI, Moscow, Russia; Sber AI Lab, Moscow, Russia; AI center block Services, Sber, Moscow, Russia; Federal Research Centre of Epidemiology and Microbiology named after Honorary Academician N. F. Gamaleya, Ministry of Health, Moscow, Russia; Laboratory of Computational Proteomics, Institute of Cytology and Genetics SB RAS, Novosibirsk, Russia

## Abstract

One of the primary tasks in vaccine design and development of immunotherapeutic drugs is to predict conformational B-cell epitopes corresponding to primary antibody binding sites within the antigen tertiary structure. To date, multiple approaches have been developed to address this issue. However, for a wide range of antigens their accuracy is limited. In this paper, we applied the transfer learning approach using pretrained deep learning models to develop a model that predicts conformational B-cell epitopes based on the primary antigen sequence and tertiary structure. A pretrained protein language model, ESM-1b, and an inverse folding model, ESM-IF1, were fine-tuned to quantitatively predict antibody-antigen interaction features and distinguish between epitope and non-epitope residues. The resulting model called SEMA demonstrated the best performance on an independent test set with ROC AUC of 0.76 compared to peer-reviewed tools. We show that SEMA can quantitatively rank the immunodominant regions within the RBD domain of SARS-CoV-2. SEMA is available at https://github.com/AIRI-Institute/SEMAi and the web-interface http://sema.airi.net.

## 1 Introduction

Selection of B-cell antibodies specifically targeting the external antigen proteins is a natural immune response *in vivo*. Corresponding antibody binding sites are called conformational B-cell epitopes, and their knowledge is important for the effective design of peptide- and protein-based vaccines and development of immunotherapeutic drugs (Gershoni et al., 2007). To date, multiple methods have been developed using machine learning and other approaches to predict conformational B-cell epitopes within an antigen sequence. For example, widely used tools include SEPPA3, Bepipred2.0, PEPITO, Epitopa, DiscoTope (Zhou et al., 2019; Jespersen et al., 2017; Sweredoski and Baldi, 2008; Rubinstein et al., 2009; Kringelum et al., 2012). However, improving the accuracy of prediction of conformational B-cell epitopes is still of great importance.

Deep learning approaches are applied increasingly often for the protein analysis and design. In particular, pretrained protein language models make it possible to address a wide range of protein classification tasks. The ESM-1v model is one of the largest transformers-based protein language models trained in a self-supervised fashion (Rives et al., 2021). Recently, the ESM-IF1 model has been developed based on a sequence-to-sequence transformer architecture with invariant geometric input features to predict the sequence of a protein based on its tertiary fold (Hsu et al., 2022). ESM-IF1 is based on GVP-GNN (Jing et al., 2020) and generic autoregressive encoder-decoder Transformer architectures (Vaswani et al., 2017) that make it possible to solve a variety of tasks, including inverse protein folding and predicting the effect of mutations. Protein residue representation from both ESM-1v and ESM-IF1 provides a large amount of contextual information that potentially can be used to predict conformational epitopes.

In the current work, we show that prediction of B-cell conformational epitopes can be significantly improved by applying transfer learning approaches using pretrained ESM-1v and ESM-IF1 models. We fine-tuned ESM-1v and ESM-IF1 models to predict residues comprising B-cell epitopes by providing an interpretable score corresponding to the expected number of contacts of an amino acid residue with the target antibody. The best performing model was called SEMA (Spatial Epitope Modelling with Artificial Intelligence). We evaluated SEMA against an independent retrospective benchmark composed of antigen residues with no prior information on antibody binding sites before the 2020 release date. SEMA was compared with Bepipred2.0 (Jespersen et al., 2017), SEPPA3.0 (Zhou et al., 2019), PEPITO (Sweredoski and Baldi, 2008), ElliPro (Ponomarenko et al., 2008) and DiscoTope2.0 (Kringelum et al., 2012) and outperformed these tools, demonstrating the highest ROC AUC value of 0.76. The SEMA prediction score was shown to correlate with the estimated immunogenicity of epitope residues according to statistical analysis of interaction the RBD domain of SARS-CoV-2 with target antibodies. SEMA is available as an online-tool and could be used for predicting B-cell conformational epitopes.

## 2 Method

### 2.1 Benchmark generation

We generated a non-redundant conformational epitopes dataset based on the available data on antigen-antibody interacting complexes in the PDB database. The pipeline used to generate the conformational epitopes dataset included the following steps:

1. The ANARCI tools was used to screen sequences of protein structures published in the PDB database that comprise heavy and light chains of Fabs (Dunbar and Deane, 2016).
2. Heavy/light Fab pairings were identified by calculating the distances between subunit residues corresponding to heavy and light chains, and only heavy and light chains with direct contacts of non CDR regions within distance of 4.5 Å were considered as the heavy/light pair. CDR loops were defined using Clothia numbering based on annotation by the ANARCI tool. Identified fab pairs were manually inspected to filter out artefacts.
3. Protein subunits that were not annotated as an antibody and had at least 5 residues interacting with antibody residues within a radius of 4.5 Å with L1/L2/L3 or H1/H2/H3 CDR loops of antibodies were considered as an antigens.
4. The contact number was calculated as the number of interactions of antigen residue atoms with antibody atoms within the distance radius of *R*. For each antigen residue, we considered two options for estimating the contact number based on the selected radius *R*: in the first case we calculated number of atoms of antibody residues in contact with any atom of antigen residue_*i*_ within the distance radius *R* (cn_atom). In the second case, we calculated the number of antibody residues in contact with any atom of antigen residue_*i*_ within the distance radius *R* (cn_aa).
5. The calculated contact numbers were mapped on the full primary sequence of the antigen. The full sequence was extracted from PDBSeqres records. All residues missing in the protein tertiary structure but present in the primary sequence were labelled as “unknown”. “Unknown” residues were excluded during model training.
6. To avoid redundancy, antigen sequences were clustered according to the degree of sequence identity (*>* 95%) using MMseqs2 software (Mirdita et al., 2019). MMseqs2 enables sensitive protein sequence searching for the analysis of massive data sets. Sequences from the same cluster were aligned using MAFFT (Katoh and Standley, 2013) and consensus epitope labels were assigned for the center of the cluster. For each residue in the reference sequence, the consensus contact number value was assigned as the maximum contact number observed among antigen-antibody complexes in the PDB dataset within an identity cluster.
7. Each antigen residue was assigned one of the following three labels. The “epitope” label was assigned according to the distance radius R1 within which at least one Fab pair residue was observed. The following values of *R*1 were considered: 4.5, 6.0 and 8.0 Å. The “close” label was assigned when the antigen residue was located outside *R*1 but within the radius *R*2 (*R*2 > *R*1) of the closest antibody residue. The following values of *R*2 were considered: 12.0, 14.0 and 16.0 Å or infinite. Remaining antigen residues were labeled “distant”. Both “close” and “distant” residues were considered as non-epitope residues.
8. To fairly compare the precision of the SEMA with other tools, we generated a retrospective data set. The data set was split into training and test sets according to the structure release date. The test set included structures that were first released in the PDB database after January 1, 2020 with no available homologous with a degree of sequence identity of 70% before this date. The remaining antigens were divided into training and validation sets in a ratio of 9:1.

### 2.2 ESM-1v and ESM-IF1 fine-tuning

ESM-1v and ESM-IF1 were fine-tuned independently by adding fully-connected liner layer after the last layer of pretrained models having an embeddings size of 1280 and 512 respectively. Outputs of the linear layer were used to predict the log-scaled contact number values for each antigen residue with the corresponding position.

Log-scaled contact number vectors were used as a target for the model. We used the Adam optimizer and the masked mean squared error loss defined as a mean squared error loss function ignoring masked residues labeled as “distant” and “unknown” residues class.

The ESM-1v model was trained for two epochs with a starting learning rate of 1e−5 and linear learning rate decay. The ESM-IF1 model was trained for two epochs with a starting learning rate of 1e−4 and linear learning rate decay.

The final models were obtained as an ensemble of five models fine-tuned independently from the same pretrained checkpoint.

As a limitation, the original ESM-1v model was pretrained with a maximum sequence length of 1022. Accordingly, sequences longer than 1022 residues were trimmed for C-terminal to the length of 1022.

### 2.3 Prediction using peer methods

Protein sequences of the tested antigen were submitted to the Bepipred2.0 server (https://services.healthtech.dtu.dk/service.php?BepiPred-2.0), and the results were downloaded in csv format (50 per run). Discotope has a stand-alone implementation on Python and was run on our own server (Kringelum et al., 2012). ElliPro has stand-alone implementation on java and was run on our own server (Ponomarenko et al., 2008). Antigen structures were submitted to the BePro server, also known as PEPITO (http://pepito.proteomics.ics.uci.edu/). The same PDB identifiers and chains were selected for submission to the SEPPA3.0 prediction server (http://lifecenter.sgst.cn/seppa/index.php) and score files were retrieved and used for metrics evaluation. All tools were run with default options.

## 3 Results

### 3.1 Benchmark for predicting epitope features

Crystallographic data on antigen-antibody structures are commonly used to identify conformational B-cell epitopes. In this paper, we screened the PDB database to select the antigen epitope residues interacting with the antibody. For each antigen residue, we calculated the contact number feature, which indicates the number of contacts of the antigen residue with antibody residues within distance radius *R*1. The resulting benchmark generated using the pipeline (see Methods) contained a total of 4,739 records, with 884 antigen sequences clustered based on the degree of identity of 95%. The test set included 101 antigen sequences.

Antigen residues were considered as epitope if the distance to the interacting antibody was lower than specified cut-off value (*R*1). *R*1 was selected in the range 4.5, 6.0 and 8.0 Å. The cut-off value of 4.5 Å reflects the presence of direct interaction with antibody residues. Radius values of 6.0 Å and 8.0 Å additionally include residues involved in long-range interaction. It is well known that epitopes can be spatially distributed on the antigen structure and for some cases such experimental information might be missing. To consider this, we split non-epitope residues based on the distance from the interacting antibody (*R*2) into “close” (*R* < *R*2) and “distant” (*R* > *R*2) (Figure 1). We selected *R*2 equal to either 12.0, 14.0 or 16.0 Å to analyze the effect of epitope boundary region information on model accuracy.

**Fig. 1.**
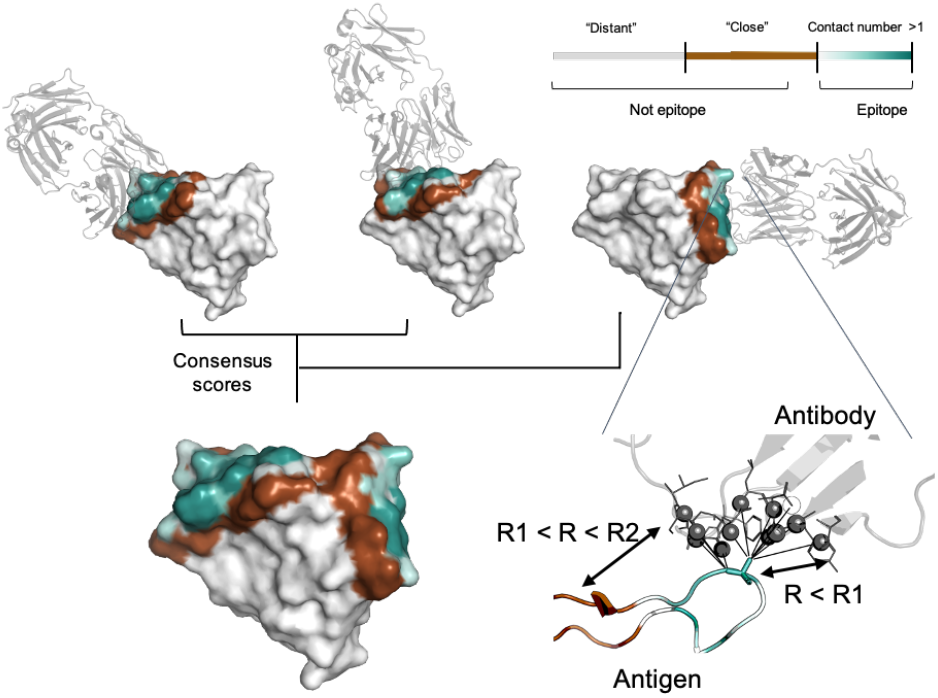
Epitopes data set generation. For each residue, the contact number interaction feature was calculated, which corresponds to log-scaled number of contacts of antigen residues with antibody residues within a radius *R*1. Antigen residues within *R*1 and *R*2 distance of the antibody were classified as non-epitope (brown color). Residues located further than *R*2 were either considered as non-epitope (the “unmasked” data set) or ignored in the model training and and calculation of the relevant metrics (the “masked” data set, highlighted in gray). The color gradient indicates the contact number value ranked from low to high. The color map is shown in the top right-hand corner of the figure. Epitopes obtained from distinct antigen-antibody complexes from the PDB database were merged to provide the final antigen epitopes data set.

In addition to conventional classification of epitope residues, for each antigen residue we calculated the contact number interaction feature. The contact number is a measure of the number of contacts of an antigen residue with atoms of antibody residues. Contact numbers may provide an additional interpretive score reflecting how deeply the residue is buried on the antigen/antibody interface. This might improve training efficiency by providing additional spatial information to the model. Finally, we combined information of different antibody for the same antigen into consensus mask. The summary of the generated consensus mask is shown in Figure 1.

This data set was used to train and evaluate the models to solve following tasks: (1) the conventional task of binary classification of antigen residues into epitope/non-epitope residues (with both “close” and “distant” residues classed as “non-epitope”); (2) prediction of epitope residues on the “masked” data set, that includes only “epitope” residues and residues localized “close” to epitope, excluding “distant” residues from model training and metrics calculation; (3) quantitative prediction of contact number features of antigen residues. We suggest that evaluation of the model on the different data sets generated using a wide range of *R*1 and *R*2 radii allow to evaluate the robustness of the model in terms of predicting epitope residues independent of the ambiguities in the epitope definition.

### 3.2 Fine-tuning the models and internal validation of SEMA

The generated conformational epitopes data set was used to fine-tune pretrained ESM-1v and ESM-IF1 models to predict the contact number antigen-antibody interaction features together with binary classification into epitope/non-epitope residues. ROC AUC metric was taken to estimate model performance. We analyzed models performance for two groups of test sets: (1) “unmasked” test sets, where all antigen residues were classified as epitope or non-epitope according to selected radii *R*1 (4.5, 6.0 and 8.0 Å), (2) “masked” test sets, where all antigen residues located further than *R*2 (12.0, 14.0, 16.0 and infinity Å) from the antibody were “masked” and ignored in ROC AUC calculations.

Thus, we evaluated a different set of *R*1 and *R*2 radii values for training set generation to select the model that performed best on both “masked” and “unmasked” test sets independent of selected radii. Additionally, we evaluated two approaches to calculating the contact number: in the first case, we counted number of antibody atoms contacting antigen within the radius *R*1 (cn_atom), whereas in the second case, we calculated the number of residues with at least one contact within the radius *R*1, which resulted in a lower value (cn_aa).

For a most variants of radii the models were able to achieve a better performance when trained to predict cn_atom compared to cn_aa; this was further used for the selection of final models (Supplementary Figure S1, S2). In the case of both ESM-IF1 and ESM-1v, the highest ROC AUC values (0.77 and 0.72, respectively) were obtained for “unmasked” test sets with *R*1=4.5 Å (Figure 2, Supplementary Figure S1, S2). However, the same models results in poorer performance for “masked” test set. This might be due to the fact that classification of antigen residues located close to epitope is a more challenging task.

**Fig. 2.**
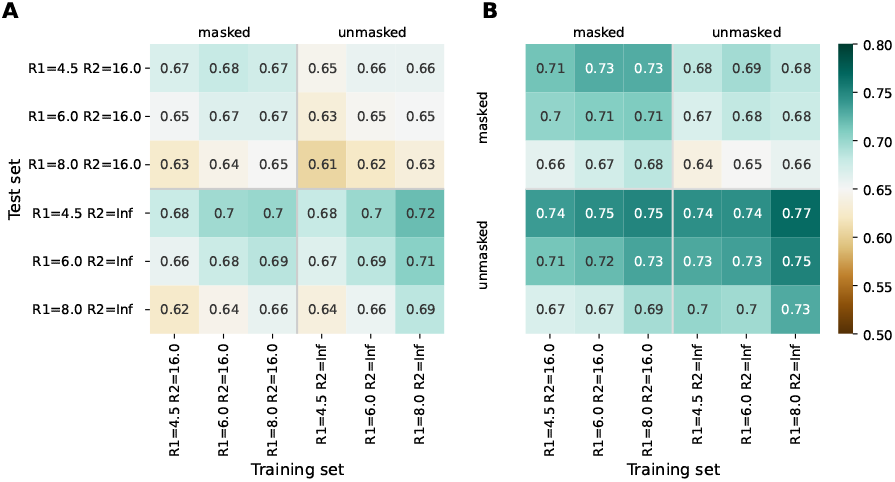
Model performance metrics estimated for the “masked” and “unmasked” test and training sets. (A) ROC AUC for SEMA-1D. (B) ROC AUC for SEMA-3D.

The models trained on *R*1 = 8.0 Å and *R*2 = 16.0 Å achieved a robust performance on all test sets independent of selected radii. Finally, ROC AUC values obtained by fine-tuned ESM-1v model was 0.7 and 0.67 for “masked” and “unmasked” test sets correspondingly (Figure 2A). ROC AUC values for ESM-IF1 fine-tuned models were 0.75 and 0.73 for “masked” and “unmasked” test set, correspondingly (Figure 2B).

The final fine-tuned models were called SEMA. SEMA involves the use of sequence-based (SEMA-1D) and structure-based (SEMA-3D) approaches to predict the conformational B-cell epitopes and provide an interpretable score indicating the log-scaled expected number of contacts with antibody residues (Figure 3).

**Fig. 3.**
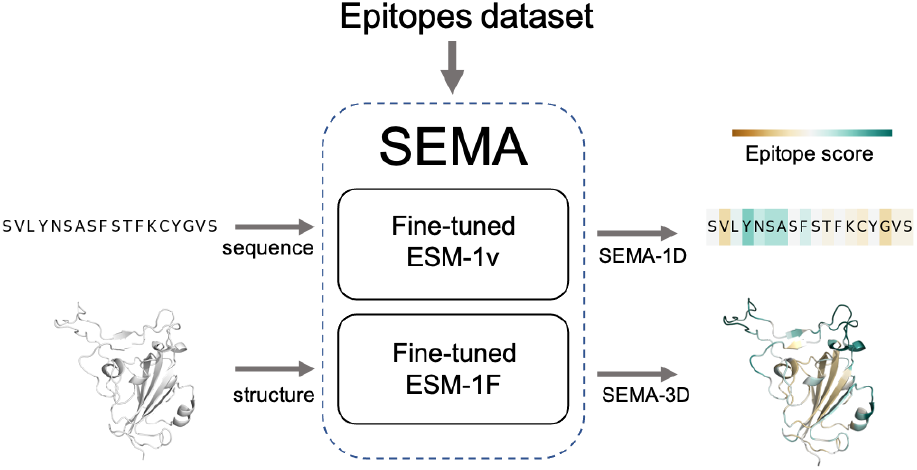
Scheme of SEMA model. SEMA comprises fine-tuned ESM-1v (SEMA-1D) for sequence-based and fine-tuned ESM-IF1 (SEMA-3D) for structure-based prediction of epitopes.

We found that calculating an ensemble of models obtained with different initialization parameters resulted in a noticeable improvement of epitope residues prediction. Thus final model was obtained as an ensemble of five fine-tuned ESM-1v models averaging their results. Fine-tuned ESM-1v models achieved best ROC AUC of 0.76/0.71 on “unmasked” and “masked” test sets respectively, whereas fine-tuned ESM-IF1 models achieved achieved best ROC AUC of 0.76/0.73 (Figure 4).

**Fig. 4.**
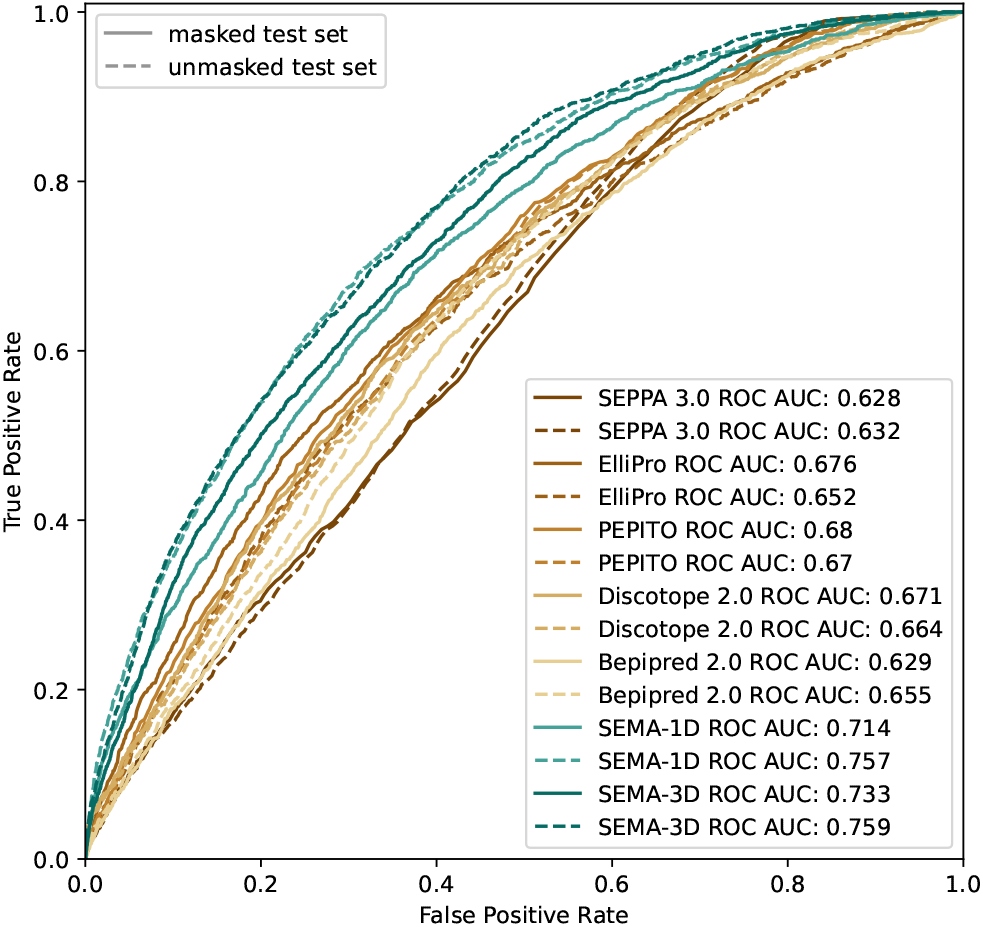
ROC AUC metrics for peer methods, SEMA-1D and SEMA-3D for two reference test sets: “masked” (*R*1 = 4.5 Å; *R*2 = 16.0 Å) and “unmasked” (*R*1 = 4.5 Å; *R*2 = *Infinite*).

### 3.3 Comparison with peer methods

The list of peer methods included Bepipred2.0 (Jespersen et al., 2017), Discotope (Kringelum et al., 2012), PEPITO (Sweredoski and Baldi, 2008), Epitopia (Rubinstein et al., 2009) and SEPPA 3.0 (Zhou et al., 2019). Model performance was evaluated against collected benchmarks (see Methods). Benchmarks included antigens with no prior information in the PDB database before January 1, 2020. These cases were not included in any training set, which enabled a fair comparison of tools with each other.

We compared the AUC metrics both for “masked” and “unmasked” test sets (Figure 4). For the test set classification into epitope and non-epitope residues, *R*1 was set equal to 4.5 Å, which was also used in the training set for other tools. In the masked test set, *R*2 was 16.0 Å.

The results shows that sequence-based methods, as well as SEPPA 3.0 and SEMA-3D better perform in conventional epitope prediction task on “unmasked” test set compared to “masked” test set. This indicates poorer performance in classification of non-epitope residues located close to epitope and predicting epitope borders. In contrast, PEPITO, ElliPro and Discotope 2.0 tools demonstrated the highest ROC AUC value on the “masked” test set. Compared to other methods, both SEMA-1D and SEMA-3D models have the highest AUC metric in all benchmark tasks. The results of models prediction for other sets of *R*1 and *R*2 radii show a similar trend and are shown in Supplementary table (see Supplementary Table S1, S2). As expected in line with SEMA, all tools had lower AUC values for higher *R*1 radii values, while selecting finite *R*2 (*R*2 < *Inf*) radius doesn’t significantly affect results.

### 3.4 Case study: Prediction of immunodominant regions of the SARS-CoV-2 RBD domain

The RBD domain of the S-protein of SARS-CoV-2 is one of the most well characterized antigens to date in terms of structure. We analyzed on the RBD domain instead of full-length S-protein to exclude the putative effect of glycosylation that is currently not considered in the SEMA (Reis et al., 2021). To evaluate the performance of SEMA, during model training we excluded all homologous sequences of the S-protein (with a degree of identity of > 70%), in particular the S-proteins of MERS and SARS-CoV. SEMA-3D was evaluated to addressed three issue: (1) correctly assigning epitope and non-epitope residues; (2) correctly predicting the contact number features; (3) predicting the immunodominant epitope residues. The immunodominant residues of RBD were estimated according to the ratio of RBD/antibody complexes in the PDB database in which RBD residue was in direct contact with antibody. We hypothesize that the calculated ratio allow to estimate the immunogenicity of RBD residues, with a high ratio corresponding to immunodominant residues.

As shown in the Figure 5, SEMA-3D provides high correlation coefficients for both contact number values and estimated immunogenicity score. Additionally, we calculated the ROC AUC metrics of the model to differentiate immunodominant residues (high ratio) from other residues (low ratio), based on the ratio threshold. This provides a more reliable estimation of model performance since most of the solvent-exposed residues of the RBD domain are labeled as epitope due to the presence of at least one structure where corresponding residues interact with the antibody. As can be seen from the score cut-off values, SEMA-3D achieves average ROC AUC metric of 0.75 on this task.

**Fig. 5.**
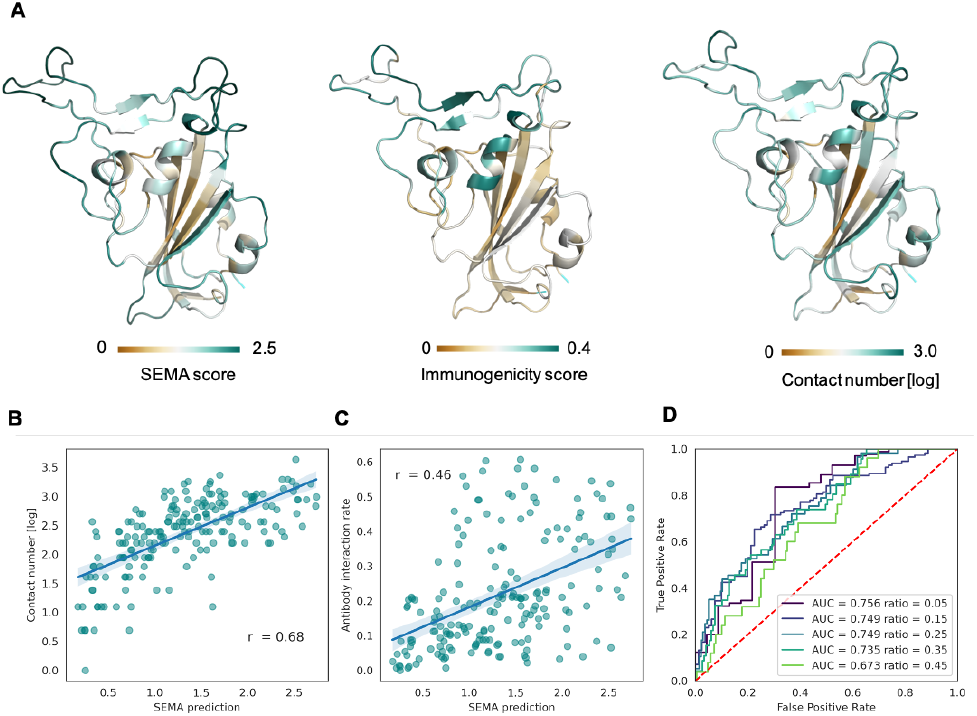
Prediction of RBD immunodominant epitopes with SEMA. ((A)) RBD domain of SARS-CoV-2 (PDB ID 7KS9, chain B) colored according to the SEMA predicted score (left), immunogenicity score (center), contact number values (right). Residues colored from brown (low value) to cyan (high value). Immunogenicity was estimated as the ratio of RBD/antibody complexes in the PDB database in which RBD residue was in contact with antibody within 8.0 Å. (B) Correlation between the SEMA score and log-scaled antigen contact number feature. Pearson correlation coefficient is shown. (C) Correlation between the SEMA score and the immunogenicity score. Pearson correlation coefficient is shown. (D) ROC AUC values calculated for different epitope/non-epitope residue classification based on the immunogenicity score threshold. ROC AUC values and threshold values for classification are denoted.)

### 3.5 Web-interface/Usage

We developed a web-interface (http://sema.airi.net) for convenient usage of SEMA. A user could either submit a protein sequence to run the fine-tuned ESM-1v model (SEMA-1D) or a protein structure in the PDB format to run the fine-tuned ESM-IF1 model (SEMA-3D). The output includes predicted epitope scores for each residue in the protein sequence. To visualize the results, the output sequence in the web-interface is colored based on the predicted contact number, with colors ranging from brown (non-epitope) to cyan (epitope) (Figure 6). In case of SEMA-3D, output includes 3D structure of protein colored using the same color scheme as in SEMA-1D (Figure 6). A user can download the results in JSON and CSV format. We also provided a code of the model implemented as the Jupyter Notebook on GitHub and available via link https://github.com/AIRI-Institute/SEMAi. We recommended using this implementation for comprehensive analysis including multiple protein sequences.

**Fig. 6.**
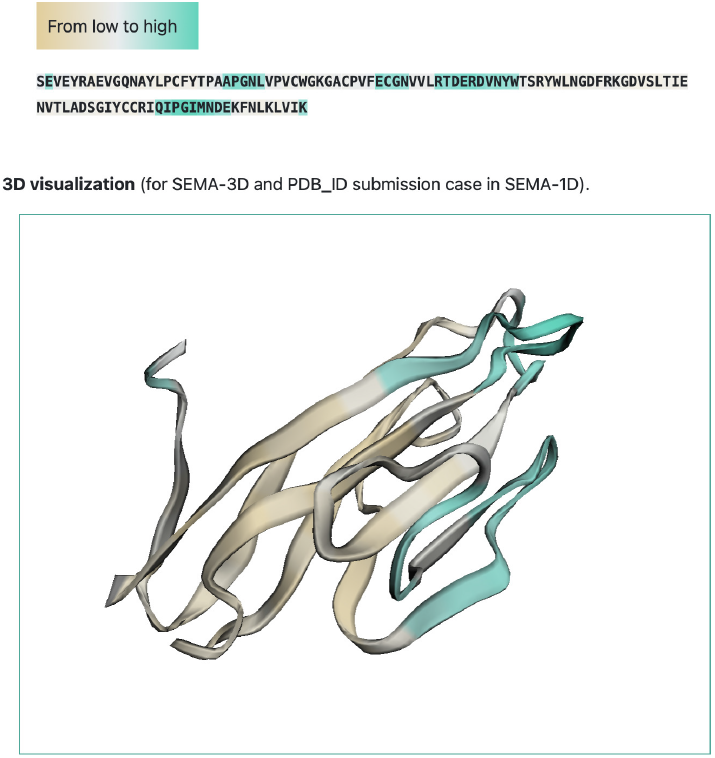
Example of SEMA graphical output (PDB ID: 6TXZ_D). Residues are colored from brown (non-epitope) to cyan (epitope).

## 4 Discussion

Computational prediction of conformational epitopes is of high importance for vaccine design and therapeutics development. However, the development of high-accuracy prediction tools is a challenging task, in particular due to the limited amount of experimental data and uncertainties in defining epitope residues. Conformational epitope residues are normally defined by distance cut-off radius between antigen and antibody residues in the interacting complex. However, arbitrary choice of distance cut-off radius might lead to ambiguous epitope label assignment.

Conventionally, a residue is classified as non-epitope if it has no interaction with the antigen. However, if experimental data on the analyzed antigen are limited, negative labels might be assigned incorrectly. In particular, for the S-protein of SARS-CoV-2, first epitopes were crystallographically discovered against the RBD domain (Yuan et al., 2020), but later immunodominant epitopes within the NTD-domain and other regions of the S-protein were also identified (Cerutti et al., 2021).

To take this problems into account, we generated a benchmark that included antigens with classified epitope residues based on two distance cut-off values. The first distance, *R*1, defined the positive epitope label class, while the second distance, *R*2, defines if the residue is too remote from the epitope and was ignored in metric calculations. Finite *R*2 radii make it possible to evaluate the model’s ability to predict the boundaries of epitopes. Additionally, for each antigen residue we calculated the contact number feature corresponding to the number of atoms of the antibody located within the radius *R*1 of antigen residue. This feature was introduced for model training providing additional spatial information on interaction between the antibody and antigen. Moreover, this feature alone was demonstrated to be a good predictor of the epitope residue for a wide range of *R*1 values.

Transfer learning was proved to be an efficient approach in the case of a limited set of examples (Howard and Ruder, 2018). In this paper, we show that a fine-tuned protein language model (ESM-1v) and an inverse folding model (ESM-IF1) perform well when predicting conformational epitopes. More specifically, the model was fine-tuned on a non-redundant set of only 783 antigen records with epitope residues assigned according to available antigen/antibody structures in the PDB database and selected *R*1 and *R*2 radius values.

To fine-tune the model, we screened the different training sets generated according for a wide range of *R*1 and *R*2 radius values and selected the model that performed well in all benchmark tasks. The final model was called SEMA; it comprises SEMA-1D (fine-tuned ESM-1v) and SEMA-3D (fine-tuned ESM-IF1) models for sequence-based and structure-based conformational B-cell epitopes prediction respectively. SEMA achieved high metrics for all benchmark tasks and was trained on the masked data set with *R*1 = 8.0 Å and *R*2 = 16.0 Å.

Additionally, we show that SEMA can predict immunogenicity of RBD domain residues. In this case we evaluated immunogenicity of the RBD domain residues as a ratio of complexes in which corresponding residue is in direct contact with antibody among all available RBD/antibody complexes.

## Author Contributions

TIS and NIV did dataset generation and analysis. TIS, NIV, IL did model training. DU wrote code for model fine-tuning. TIS and AVK did analysis of published epitope tools performance. DNS, MSA, PVS, TIS and NIV did manuscript writing. MS and TIS developed web-interface. OLK, DNS, MSA, NIV designed the study.

## Acknowledgments

This study was performed using the infrastructure of AIRI, Artificial Intelligence Research Institute (Moscow, Russia). Conflicts of interest: none declared.

## Data Availability

The datasets generated for this study can be found in the GitHub repository: https://github.com/AIRI-Institute/SEMAi.

## References

Cerutti, G., Guo, Y., Zhou, T., Gorman, J., Lee, M., Rapp, M., et al. (2021). Potent sars-cov-2 neutralizing antibodies directed against spike n-terminal domain target a single supersite. Cell host & microbe 29, 819–833

Dunbar, J. and Deane, C. M. (2016). Anarci: antigen receptor numbering and receptor classification. Bioinformatics 32, 298–300

Gershoni, J. M., Roitburd-Berman, A., Siman-Tov, D. D., Freund, N. T., and Weiss, Y. (2007). Epitope mapping. BioDrugs 21, 145–156

Howard, J. and Ruder, S. (2018). Universal language model fine-tuning for text classification. arXiv preprint 1801.06146

Hsu, C., Verkuil, R., Liu, J., Lin, Z., Hie, B., Sercu, T., et al. (2022). Learning inverse folding from millions of predicted structures. bioRxiv

Jespersen, M. C., Peters, B., Nielsen, M., and Marcatili, P. (2017). Bepipred-2.0: improving sequence-based b-cell epitope prediction using conformational epitopes. Nucleic acids research 45, W24–W29

Jing, B., Eismann, S., Suriana, P., Townshend, R. J., and Dror, R. (2020). Learning from protein structure with geometric vector perceptrons. arXiv preprint 2009.01411

Katoh, K. and Standley, D. M. (2013). Mafft multiple sequence alignment software version 7: improvements in performance and usability. Molecular biology and evolution 30, 772–780

Kringelum, J. V., Lundegaard, C., Lund, O., and Nielsen, M. (2012). Reliable b cell epitope predictions: impacts of method development and improved benchmarking. PLoS computational biology 8, e1002829

Mirdita, M., Steinegger, M., and Söding, J. (2019). Mmseqs2 desktop and local web server app for fast, interactive sequence searches. Bioinformatics 35, 2856–2858

Ponomarenko, J., Bui, H.-H., Li, W., Fusseder, N., Bourne, P. E., Sette, A., et al. (2008). Ellipro: a new structure-based tool for the prediction of antibody epitopes. BMC bioinformatics 9, 1–8

Reis, C. A., Tauber, R., and Blanchard, V. (2021). Glycosylation is a key in sars-cov-2 infection. Journal of Molecular Medicine 99, 1023–1031

Rives, A., Meier, J., Sercu, T., Goyal, S., Lin, Z., Liu, J., et al. (2021). Biological structure and function emerge from scaling unsupervised learning to 250 million protein sequences. Proceedings of the National Academy of Sciences 118

Rubinstein, N. D., Mayrose, I., Martz, E., and Pupko, T. (2009). Epitopia: a web-server for predicting b-cell epitopes. BMC bioinformatics 10, 1–6

Sweredoski, M. J. and Baldi, P. (2008). Pepito: improved discontinuous b-cell epitope prediction using multiple distance thresholds and half sphere exposure. Bioinformatics 24, 1459–1460

Vaswani, A., Shazeer, N., Parmar, N., Uszkoreit, J., Jones, L., Gomez, A. N., et al. (2017). Attention is all you need. Advances in neural information processing systems 30

Yuan, M., Wu, N. C., Zhu, X., Lee, C.-C. D., So, R. T., Lv, H., et al. (2020). A highly conserved cryptic epitope in the receptor binding domains of sars-cov-2 and sars-cov. Science 368, 630–633

Zhou, C., Chen, Z., Zhang, L., Yan, D., Mao, T., Tang, K., et al. (2019). Seppa 3.0—enhanced spatial epitope prediction enabling glycoprotein antigens. Nucleic acids research 47, W388–W394

