## Supplementary Figures 1,2 for "SEMA: Antigen B-cell conformational epitope prediction using deep transfer learning"

### Supplementary material

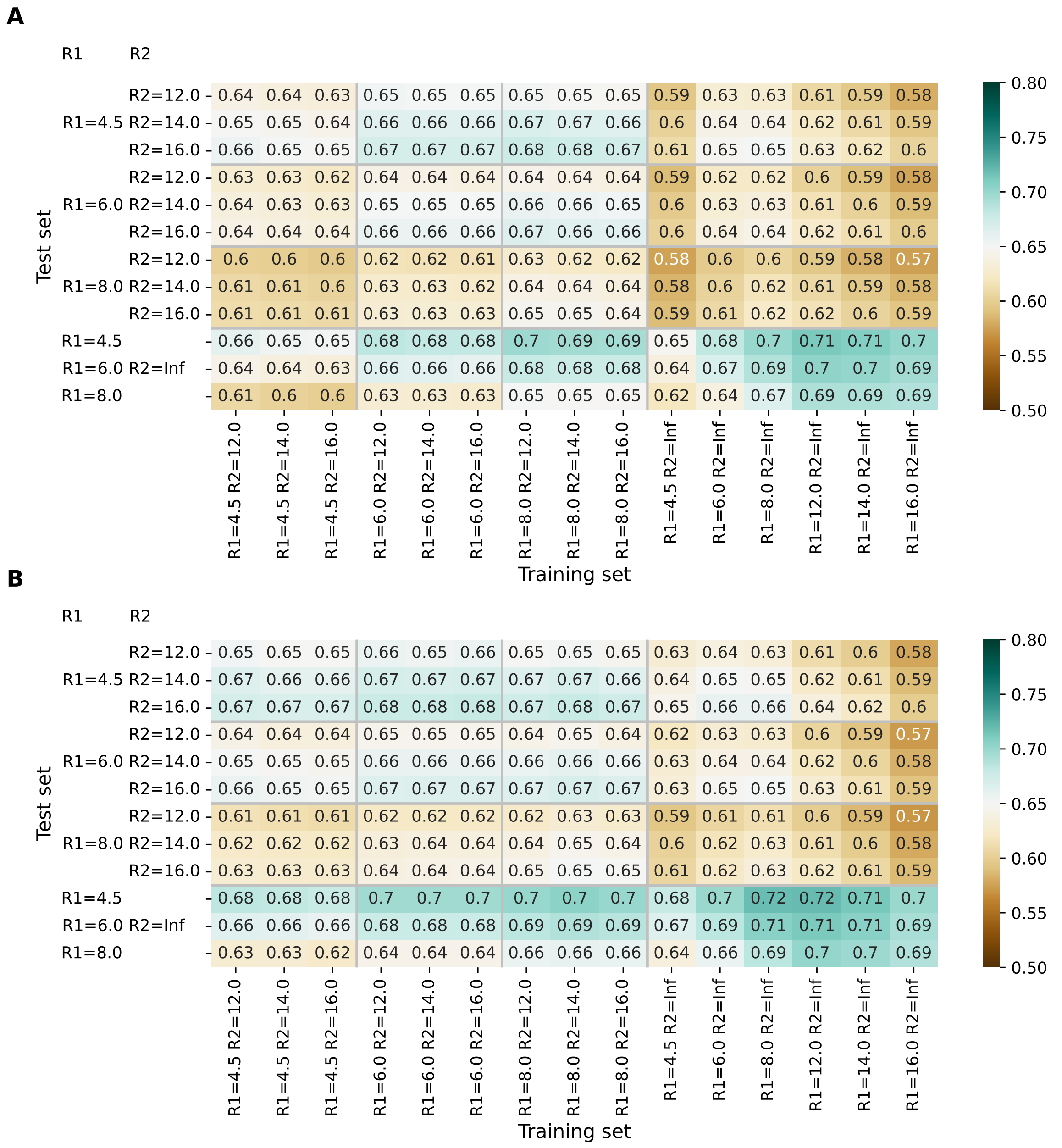


**Fig. S1.** ROC AUC values for SEMA-1D regression model for different data set. (**A**) ROC AUC values for SEMA-1D regression model based on cn_aa approach. (**B**) ROC AUC values for SEMA-1D regression model based on cn_atom approach.

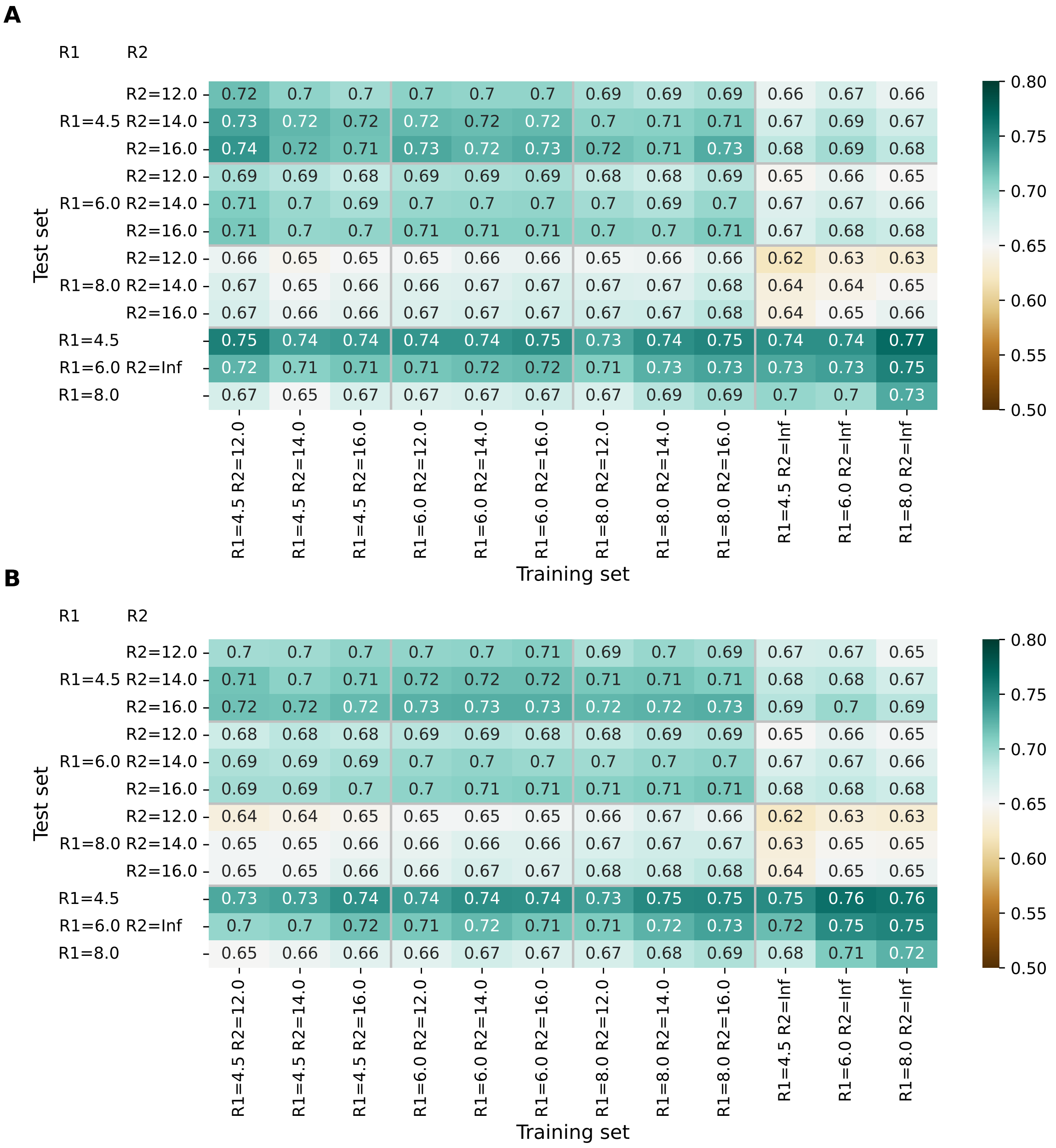


**Fig. S2.** ROC AUC values for SEMA-3D regression model for different data set. (**A**) ROC AUC values for SEMA-3D regression model based on cn_aa approach. (**B**) ROC AUC values for SEMA-3D regression model based on cn_atom approach.
